# Characterization of kinase gene expression and splicing profile in prostate cancer with RNA-Seq data

**DOI:** 10.1101/061085

**Authors:** Huijuan Feng, Tingting Li, Xuegong Zhang

**Author notes:** Email addresses: Huijuan Feng.

## Abstract

**Background**

Alternative splicing is a ubiquitous post-transcriptional process in most eukaryotic genes. Aberrant splicing isoforms and abnormal isoform ratios can contribute to cancer development. Kinase genes are key regulators of various cellular processes. Many kinases are found to be oncogenic and have been intensively investigated in the study of cancer and drugs. RNA-Seq provides a powerful technology for genome-wide study of alternative splicing in cancer besides the conventional gene expression profiling. But this potential has not been fully demonstrated yet.

**Methods**

Here we characterized the transcriptome profile of prostate cancer using RNA-Seq data from viewpoints of both differential expression and differential splicing, with an emphasis on kinase genes and their splicing variations. We built up a pipeline to conduct differential expression and differential splicing analysis. Further functional enrichment analysis was performed to explore functional interpretation of the genes. With focus on kinase genes, we performed kinase domain analysis to identify the functionally important candidate kinase gene in prostate cancer. We further calculated the expression level of isoforms to explore the function of isoform switching of kinase genes in prostate cancer.

**Results**

We identified distinct gene groups from differential expression and splicing analysis, which suggested that alternative splicing adds another level to gene expression regulation. Enriched GO terms of differentially expressed and spliced kinase genes were found to play different roles in regulation of cellular metabolism. Function analysis on differentially spliced kinase genes showed that differentially spliced exons of these genes are significantly enriched in protein kinase domains. Among them, we found that gene *CDK5* has isoform switching between prostate cancer and benign tissues, which may affect cancer development by changing androgen receptor (AR) phosphorylation. The observation was validated in another RNA-Seq dataset of prostate cancer cell lines.

**Conclusions**

Our work characterized the expression and splicing profile of kinase genes in prostate cancer and proposed a hypothetical model on isoform switching of *CDK5* and *AR* phosphorylation in prostate cancer. These findings bring new understanding to the role of alternatively spliced kinases in prostate cancer and demonstrate the use of RNA-Seq data in studying alternative splicing in cancer.

## Background

Alternative splicing is an important post-transcriptional regulation through which one gene can produce multiple isoforms. Alternative splicing is found to be ubiquitous in human cells, and about 95% of human multi-exon genes undergo this process [1]. The multiple protein isoforms generated with this mechanism play critical roles in diverse cellular processes such as cell cycle control, differentiation and cell signaling [2]. Aberrant alternative splicing has been reported to be highly relevant to many human diseases including cancer. People have found that many cancer-related genes undergo alternative splicing and cancer-specific alternative splicing events contribute to carcinogenesis [3]. RNA-Seq technology and the bioinformatics methods have provided great opportunities for studying alternative splicing in cancer [4].

Protein kinases are one of the largest gene families in human and they constitute ~1.7% of human genes [5]. Protein kinases are key regulators of cell functions. They can mediate important cellular processes like signal transduction and cell cycle. Kinases regulate the activity, localization and function of substrate proteins by adding phosphate groups to them [5]. Mutations and dysregulation of protein kinases have been found to be causal reasons of some human diseases especially cancer. Numerous efforts have been put to target oncogenic kinases by developing inhibitors for disease therapy [6]. Merkin et al. have revealed that a large percentage of alternative splicing events often contribute to alterations in protein phosphorylation and kinase signaling [7]. It indicated the high relevance of alternative splicing with protein phosphorylation. It has been observed that some aberrant splicing events can modify kinase activities by truncating the kinase domain or fine-tuning the binding specificity to functional partners [8].

Prostate cancer is a major type of cancer in men. Normal prostate tissue development needs steady activity of androgens with androgen receptors (*AR*). Androgens like testosterone and dihydrotestosterone (DHT) can bind to *AR* and induce *AR* transcriptional activity. *AR* is a member of the steroid receptor superfamily and also a ligand-activated nuclear transcription factor. Most prostate cancers are dependent on the androgen-*AR* interaction for cell growth and proliferation initially. Androgen ablation therapy can lead androgen-dependent cancer to be repressed [9], but some of them cannot be cured and become androgen-independent in a hormone refractory state. Modulation of *AR* transcriptional activity is conducted by interactions of *AR* and co-regulators. Some signal transduction pathways with various kinases involved can also regulate *AR* transcriptional activities via phosphorylation of *AR* and *AR* co-regulators, and may be a main approach to maintain *AR* and co-regulator transcriptional activity in *AR*-negative prostate cancer cells [10]. Moreover, kinases play key roles in cell proliferation and cell homeostasis maintenance, and the deregulation of kinases related to important signal transduction pathways can lead to initiation and progression of cancer [11]. It’s important to investigate the expression profile of kinases to explore those that take parts in the pathogenesis of prostate cancer, especially those involved in *AR* activity.

Many efforts have been devoted to study kinases that are deregulated in cancer and may serve as potential targets for cancer treatment. Differentially expression of kinase genes in prostate cancer has been studied with RT-PCR [12] and microarrays [13], but the alternative splicing of these genes have not been systematically studied in prostate cancer. The inclusion/exclusion of exons by alternative splicing can produce multiple isoforms that can vary in protein structure and functions. Abnormal isoform variants and abnormal isoform expression ratios can lead to dysregulation of various cancer-related genes and pathways [3]. It’s important to study differentially spliced kinases that may lead to functional variations in kinase activities in prostate cancer.

In this study, using an RNA-Seq dataset published by K. Kannan et al. [14], we characterized the transcriptome profile in prostate cancer from viewpoints of both differential expression and differential splicing, with an emphasis on kinase genes and their splicing isoforms. We profiled genes that are differentially expressed (DE) and differentially spliced (DS) between prostate cancer and benign tissues. GO and KEGG pathway enrichment analyses revealed distinct functions in prostate cancer. We took further study on the kinases among the detected genes and identified kinases that may have function alterations by differential splicing in prostate cancer through protein domain analysis. Among them, the *CDK5* was detected to undergo isoform switching in prostate cancer and benign tissues, which suggests an important regulatory role of *CDK5* on *AR* phosphorylation in prostate cancer via alternative splicing. The result was validated on another RNA-Seq dataset of prostate cancer cell lines[15], and also has implications in difference between androgen-dependent and androgen-independent cancer progression.

## Methods

To characterize the transcriptome profile in prostate cancer, we analyzed a RNA-Seq dataset by K. Kannan et al. [14] using the strategy we discussed in [4]. The pipeline of the analysis are shown in Figure 1, with details provided in the sections below.

**Figure 1.**
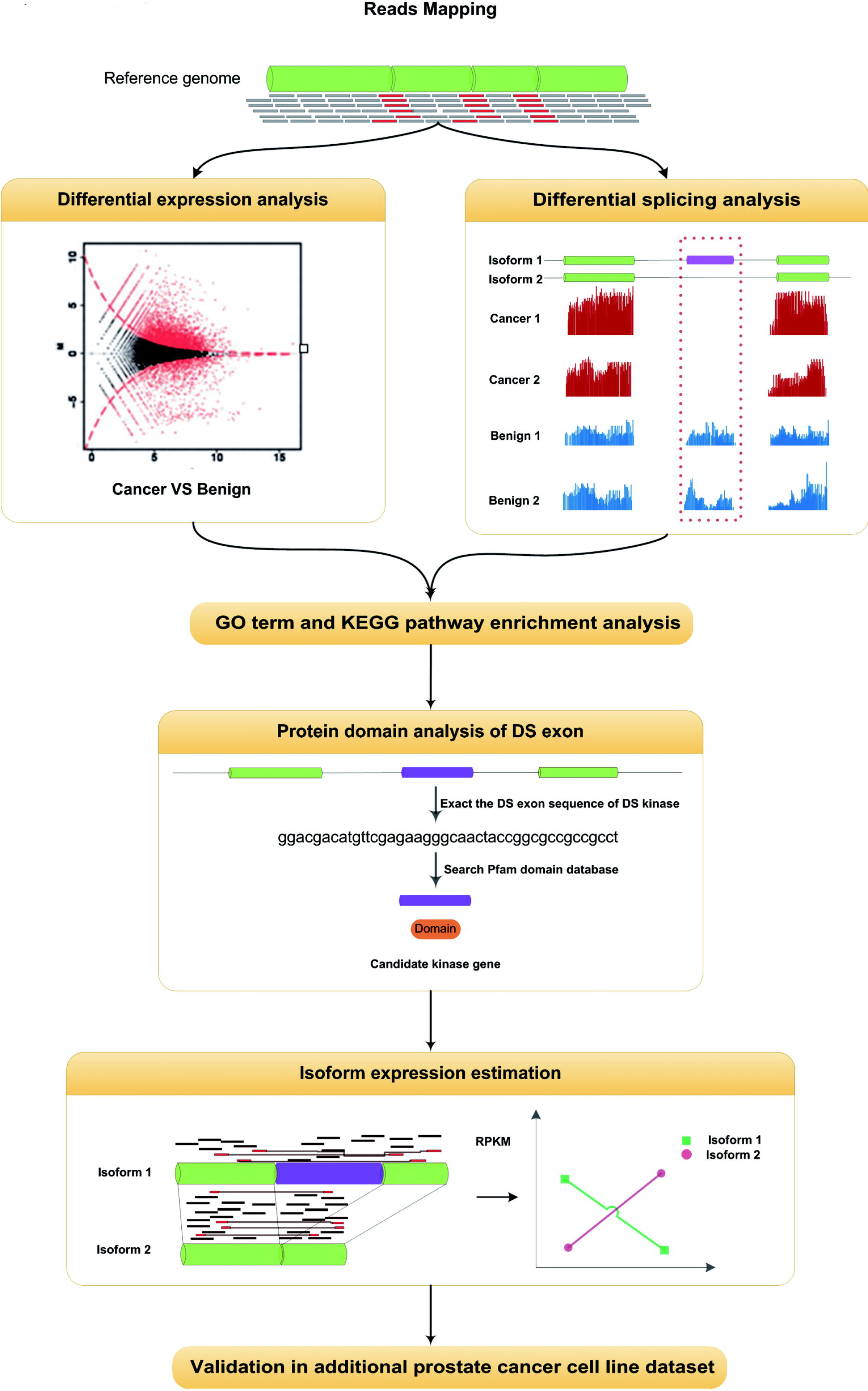
Overview of data analysis pipeline. Reads were first mapped to human reference genome. Differential expression and differential splicing analysis between cancer and benign samples were followed with a special focus on kinase genes. GO and KEGG pathway enrichment analysis were performed on the DE and DS genes to explore functions of them. Pfam domains were searched for the DS kinase exons to identify the candidate kinases whose splicing alternations are functionally important in prostate cancer. Isoform expression analysis was conducted to confirm the isoform switching of candidate DS kinase genes. The candidate DS kinase results were validated on additional prostate cancer cell line data.

### Data

The RNA-Seq data of 20 prostate cancer and 10 matched benign tissue samples [14] were downloaded from NCBI SRA database (accession number SRP002628). The data were obtained by 36bp paired-end transcriptome sequencing with Illumina GAII. The prostate cancer cell line dataset was obtained from SRA (accession number SRP004637) generated by John et al. [15]. The dataset contains 21 prostate cell lines sequenced by Illumina GA II. Cell line samples have various number of replicates. The prostate cancer cell lines includes benign, androgen-dependent and androgen-independent cell lines.

### Reads mapping

The short sequencing reads were first remapped to human reference genome using TopHat (version 1.4.1) [16]. Uniquely mapped reads with no more than two mismatches were kept for downstream analysis. Bam files generated by TopHat using 17 as the “–segment-length” parameter were first converted to bed format files for the differential expression analysis.

### Differential expression analysis

Identification of DE genes was conducted using DEGseq (version 1.2.2) [17]. An MA-plot-based method was used to identify DE genes between the groups of the 20 prostate cancer samples and the 10 prostate benign samples using normalized gene expression values (RPKM) as input. Genes were reported as differentially expressed for downstream analysis at two significance cut-offs:p-value < 0.05 and FDR < 1%.

### Differential splicing analysis

We used DSGseq [18] and DEXSeq (version 1.6.0) [19] to detect DS genes between the two groups. DSGseq uses negative-binomial distribution to model read counts on exons and defines NB-statistic to detect difference in splicing of exons between two groups. The exon information was from hg18 annotation, and bam format mapping files were given to DSGseq, which reported a list of DS genes along with the DS exons. The cut-off of NB_stat=2 was used, corresponding to a moderate stringency. DEXSeq also detects differential splicing in an exon-centric manner with p-values reported for differential usage of each exon. It also provides visualization for differential spliced genes. We chose adjusted p-value < 0.05 as cut-off for DS exons. The consensus results of the two methods were reported as DS genes in this work. DS genes were visualized using the DEXSeq plot function.

### Gene Ontology and KEGG pathway enrichment analysis

We used GOseq (version 1.10.0) [20] to identify Gene Ontology (GO) terms which are over-represented by the detected genes. GOseq has special designs in its statistical test for avoiding possible selection bias toward long and highly expressed genes. GO terms were considered as statistically significant with false discovery rate (FDR) < 0.05. We used REVIGO [21] to eliminate redundant terms and summarize them into representative clusters. To study the pathways DE and DS kinase genes are involved in, we mapped them to KEGG pathways by the DAVID functional annotation tool [22]. Pathways with p-value < 0.05 were declared significant.

### Domain analysis of differentially spliced kinase exons

Pfam domain search [23] was carried out to analyze the protein kinase activity of the alternative splicing exons in the DS kinase genes. We built a background set by extracting all alternative exons of 518 kinase genes according to the annotation of hg18 downloaded from UCSC Genome Browser. DNA regions of both the DS exons and the background set were translated into peptides in six-frame manner using sequence translation tool “EMBOSS Transeq”. The peptides sequences were searched against Pfam database for matching domain with the default cut-off value of E-value=1.0. Fisher’s Exact Test was performed to compare the kinase domain hits of DS kinase exons and background exons.

### Isoform expression estimation

The software NURD (version 1.1.0) [24] was used to study the isoform expression levels of DS genes. It uses nonparametric models to deal with possible position-related biases in RNA-Seq data and estimates the isoform expression by maximizing a likelihood function.

## Results

### Differential expression analysis reveals gene signatures of prostate cancer

Comparing prostate cancer and benign tissue samples with DEGseq package, we identified 3,093 genes as differentially expressed with p-value < 0.05. Among them, 528 genes are differentially expressed with FDR<1% (Additional file 2). Focusing on kinase genes, we found 62 DE kinase genes at p-value< 0.05 and 16 DE kinase genes at FDR<1% (Additional file 3).

We lists the top 10 up-regulated genes and top 10 down-regulated genes (Additional file 4). Their functions related with prostate cancer by literature are also provided. Most of the top genes have been reported to be related with prostate cancer in literature. The most up-regulated gene in prostate cancer is phospholipase A2 group (*PLA2G2A*), an important enzyme in inflammatory processes. Tuomas et al. reported that the expression of *PLA2G2A* were significantly increased in prostate cancer comparing to benign tissue, and it might serve as a prognostic maker for prostate cancer [25]. The second up-regulated gene, orosomucoid 1(*ORM1*), is an androgen receptor (*AR*)-activated gene and involved in *AR* pathway[26]. Ayla et al. found that *ORM1* is differentially expressed between non-recurrent primary and metastatic prostate cancer and is involved in cancer-metabolism and immune response pathways [27]. Notably, the fifth up-regulated gene antigen 3 (*PCA3*) is a non-protein coding gene. Research on the relationship of *PCA3* and prostate cancer has been a long story since 1999 [28]. *PCA3* is prostate-specific and has been shown highly expressed in prostate cancer, and has been identified as a genetic marker for prostate cancer diagnosis [29].

The most down-regulated gene in prostate cancer is serpin peptidase inhibitor, clade A (alpha-1 antiproteinase, antitrypsin) member 3(*SERPINA3*), whose protein product, Alpha 1 antichymotrypsin (*ACT*) is associated with increased risk of prostate cancer [30]. The second down-regulated gene lactotransferrin (*LTF*) is a member of transferrin family and its protein product is a major iron-binding protein. There’s one study identified *LTF* as the most significantly down-regulated gene in prostate cancer cells and proved that *LTF* protein can inhibit the growth of prostate cancer cells. The down regulation of *LTF* might be involved in prostate cancer progression [31]. Another gene that deserves highlight is superoxide dismutase 2(*SOD2*). It encodes a mitochondrial enzyme that can protect cell from oxidative damage, and has been known as a tumor suppressor gene in human prostate cancer. Increased expression of *SOD2* has been shown to result in the suppression of prostate cancer cell growth by mediating the senescence-associated tumor suppression with insulin-like growth factor binding protein-related protein-1 (*IGFBP-rP1*) [32]. The down regulation of *SOD2* here may help promote the cancer progression. In brief, the results are consistent with previous studies (Additional file 4). Most of the top DE genes we detected are gene signatures of prostate cancer and have important functions in prostate cancer.

### Most differentially spliced genes in prostate cancer do not show differential total expression

Besides differential expression analysis at gene level, we studied possible splicing variations between cancer and benign tissues. We identified 2,651 genes that are differentially spliced between prostate cancer and benign prostate tissues (Additional file 5), including 55 DS kinase genes (Additional file 6).

We compared the profiles of DS and DE genes. The overlap of two profiles has 660 genes (Figure 2A), composing less than 25% of the DE genes. Among the 55 DS kinase genes, 12 are differentially expressed between cancer and benign samples at gene level (Figure 2B). Most of the DS kinases tend to have stable overall expression level but with different splicing isoforms and different exon usages. The small overlap between differential expression and differential splicing indicates that gene expression regulation can be modulated at both gene-level and isoform-level. Genes can have identical total expression abundances but different isoform ratios and different major isoforms, which can have significant functional impact.

**Figure 2.**
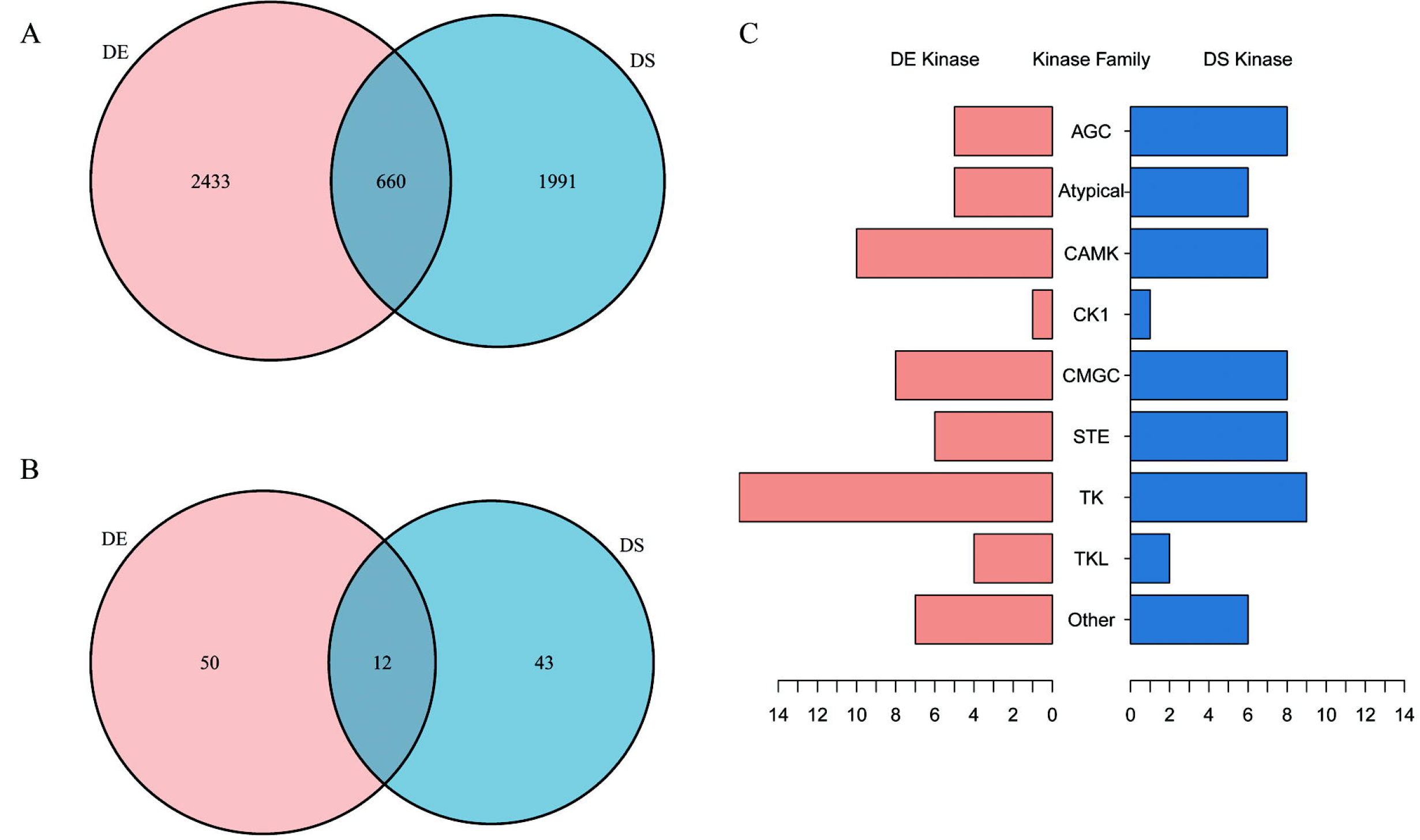
Profiles of DE and DS genes between prostate cancer and benign tissues. (A) Overlap of differential expression (DE) and differential splicing (DS) genes. (B) Overlap of DE and DS kinase genes. (C) DE and DS kinase gene counts in different protein kinase families.

The detected DE or DS kinase genes belong to 9 eukaryotic kinase families. The tyrosine kinase (TK) family has the largest number of members (Figure 2C). Tyrosine kinase can phosphorylate tyrosine amino acid on substrates specifically. TK members function in a wide variety of processes and pathways, from transmembrane signaling to signal transduction to the nucleus and cell-cycle control and transcription factor activities [33]. TK has been shown to be associated with several cancers and have implications for cancer treatment [34].

### GO and KEGG enrichment analysis show distinct functions of DE and DS genes in prostate cancer

We performed GO and KEGG pathway enrichment analysis on the detected DE and DS genes to investigate their biological relevance to prostate cancer. GO terms related with cancer and prostate cancer like cell proliferation, cell adhesion, and prostate gland development are found to be enriched by DE genes (Additional file 1 – Figure S1, Additional file 7). Enriched KEGG pathways of DE genes include ribosome, metabolic pathways, p53 signaling pathway, MAPK signaling pathway and other disease-related pathways like Parkinson’s disease, Alzheimer’s disease, pathways in cancer, prostate cancer and bladder cancer (Additional file 1- Figure S2, Additional file 7). Three genes closely related to functions of *AR* in prostate cancer, *AR, KLK3* and *MAPK3*, are also among the detected DE genes. In comparison, GO enrichment analysis of DS genes shows a wide range of GO terms related to mRNA metabolism, regulation of catalytic activity, intracellular transport, and protein complex subunit organization etc. (Additional file 1 31 – Figure S3, Additional file 7). Pathways including proteasome, ribosome, and spliceosome are found to be enriched by DS genes (Additional file 6).

We also did functional enrichment analysis of the DE and DS kinases comparing to the background of all kinases. Interestingly, enriched GO terms of DE kinases can be summarized as negative regulation of cellular metabolism, while enriched GO terms of DS kinases can be summarized as positive regulation of biological process and protein metabolic process, and both DE and DS kinases are found enriched in GO terms related to protein phosphorylation (Additional file 1- Figure S4, Additional file 7). In the KEGG pathway analysis, detected DE kinases show significant enrichment in cancer-related pathways (Figure 3A, Additional file 8), with the MAPK signaling pathway and prostate cancer at the top of the list. The MAPK pathway also comes at the top of the list of pathways enriched by DS kinase genes, followed by focal adhesion etc. (Figure 3B).

**Figure 3.**
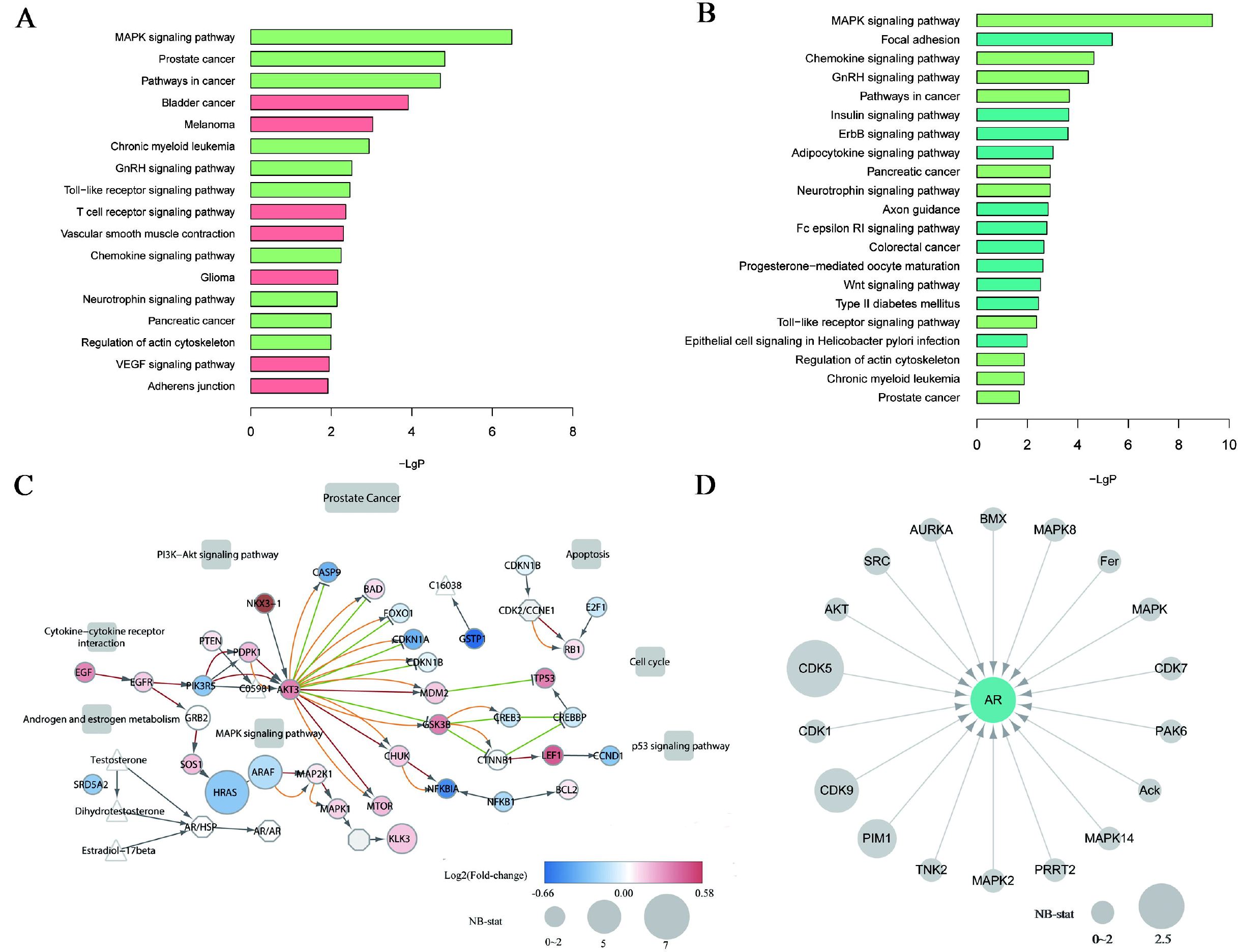
Functional enrichment analysis of DE and DS kinase genes. (A) Enriched KEGG pathways by DE kinase genes. (B) Enriched KEGG pathways by DS kinase genes. Pathways enriched in both DE and DS kinase genes are shown in green in both plots. (C) The “Prostate Cancer” pathway in KEGG, with detected DE and DS genes highlighted in colors. Circles represent genes, with their colors showing their fold-change between the two groups. Circle size indicates the NB-stat calculated by DSGseq, which represents the degree of differential splicing of the gene. Small molecules (triangles), cellular process (rectangles) and gene complexes (hexagon) are included. Edges indicate activation (red), inhibition (green), phosphorylation (orange). (D) Expression and splicing changes of kinases that phosphorylate *AR*. None of the kinase genes is differentially expressed, but *CDK5, CDK9, PIM1* and *SRC* are differentially spliced, with node sizes representing the NB-stat.

To visualize the DE and DS gene profiles in a context of pathways, we compiled the KEGG pathway map “Prostate cancer” by considering the expression and splicing difference between cancer and benign (Figure 3C). There are 8 DE genes in the pathway, including *KLK3, NKX3-1, GSTP1, NFKBIA, CCND1, CDKN1A, TP53* and *HRAS*, and there are 6 DS genes including *HRAS, ARAF, KLK3, GRB2, MDM2* and *CHUK*. The gene *KLK3* encodes the glycoprotein enzyme known as prostate-specific antigen (PSA), an important tumor marker used for diagnosis of prostate cancer in clinical practices [35]. *KLK3* is found to be both differentially expressed and spliced in the pathway. *KLK3* has 4 alternative splicing isoforms annotated in RefSeq. These differentially expressed isoforms may have different roles in prostate cancer progression. The gene *HRAS* was also found both differentially expressed and spliced. *HRAS* belongs to the *Ras* oncogene family, which encodes proteins functioning in signal transduction pathways. *HRAS* gene can produce two protein isoforms with complementary function related to cell proliferation [36]. The differential splicing of gene *HRAS* may suggest the roles of expression change at isoform level in prostate cancer. We noted that none of the kinases that can phosphorylate *AR* were differentially expressed, but 4 kinases, *CDK5, CDK9, PIM1* and *SRC* were differentially spliced (Figure 3D). Searching the literature, we found that these four DS kinases had been reported to be involved in prostate cancer in different functions [37–40].

### Differentially spliced kinase exons are significantly enriched in protein kinase domain

Most of protein kinases are involved in critical cellular activities. Splicing variability of functional region such as ATP binding domain and kinase domain may directly initiate or contribute to cancer progression. Alternative splicing may regulate oncogenic kinase activity by generating isoforms that skip kinase domain or truncate kinase domain [8]. We investigated that whether the differentially spliced exons of 55 DS kinases have functional impact on kinase activity by searching domains against Pfam database[23]. Remarkably, we found the DS exons of 16 DS kinase genes have domain hits against the Pfam domain database. Among them, 9 DS kinase genes include the protein kinase domain (Pkinase) (Table 1). To verify if the functional relevance of DS exons of the 55 DS kinases to kinase domain was more significant than all a random group of alternative splicing events of the human kinases, we collected all known alternative splicing exons of 518 kinases from UCSC Genome Browser as background and searched them in the Pfam database. In total, 81 significant Pkinase domains were detected in 8,671 peptide sequences translated from 1,936 alternative splicing exons of 518 kinases. Taking total alternative splicing events in all the kinases as background, we found that the DS exons of the 55 DS kinases are significantly related to protein kinase domain with a p-value smaller than 0.007 (Fisher’s Exact Test). Besides, we investigated that whether the kinase domain predicted by Pfam search really existed in the annotation database GeneCards [41]. We found that either truncated kinase domain or total kinase domain exist in the DS exons of 9 kinases. Especially, we found that the DS exon of *CDK5* is included in one isoform and excluded in the other. We took it as an example to conduct further isoform-level analysis.

**Table 1.** Protein kinase domain information of differentially spliced exons in kinases

| Gene name | Chr | AS_start | AS_end | Isoform including AS region | Isoform not including the AS region | Domain or active site included |
| --- | --- | --- | --- | --- | --- | --- |
| *CAMK2B* | chr7 | 44248341 | 44248464 | NM_172083,NM_172082,<br>NM_172080,NM_172078,<br>NM_172084,NM_172081,<br>NM_001220,NM_172079 |  | Truncated protein kinase domain |
| *CDK5* | chr7 | 150383772 | 150383868 | NM_004935 | NM_00116441<br>0 | Truncated protein kinase domain, proton<br>acceptor active site |
| *CHUK* | chr10 | 101970334 | 101970423 | NM_001278 |  | Inside protein kinase domain |
| *CSNK1D* | chr17 | 77804180 | 77804409 | NM_001893,NM_139062 |  | Inside protein kinase domain ,proton<br>acceptor active site |

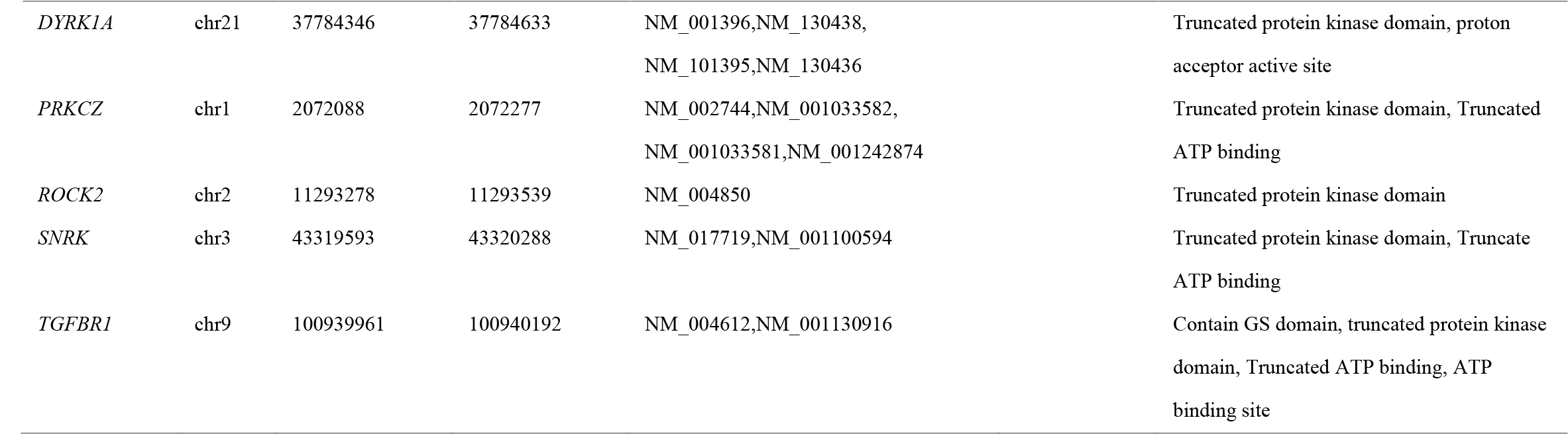

The protein functional analysis results indicated that DS exons of the DS kinases are significantly enriched in kinase domain. Differential splicing of these kinases may modify kinase activity and have functional impact on phosphorylation of their substrates in prostate cancer progression by differential expression of isoforms including or excluding those DS exons.

### Isoform switching of *CDK5* in prostate cancer and its potential functions on *AR* phosphorylation

The DS kinase gene *CDK5* has two isoforms (Figure 4A). Differential splicing analysis showed that *CDK5* is differentially spliced between prostate cancer and benign tissues (Figure 4B). The inclusion and exclusion of the 32 amino acid DS exon (exon 6) define the two isoforms. We refer the longer isoform with exon 6 to as Isoform1 (NM_00495 in REFSEQ), the shorter isoform as Isoform2 (NM_001164410).

Database searching revealed that the serine/threonine protein kinase active site lies in this alternative exon of gene *CDK5* (Additional file 1 – Figure S5). Isoform1 has full length of 293 amino acids and encodes protein *CDK5* while Isoform2 lacks exon 6 and encodes a 260 amino acids protein. *CDK5* is one of the cyclin-dependent kinase (*CDK*) family members and a serine/threonine kinase that plays important roles in various cellular activities like cell differentiation and apoptosis, etc. [42]. The function of protein encoded by Isoform2 has not been well studied in literature. Kim et al. reported different subcellular localizations of two isoforms of *CDK5* which might indicate different functions [43]. *CDK5* knockdown by siRNA resulted in changes of the microtubule cytoskeleton, loss of cellular polarity and motility in human prostate cancer DU 145 cell line [40]. It has been reported that protein *CDK5* can activate and stabilize *AR* in the nucleus through Ser-81 phosphorylation in prostate cancer cells [44]. These studies all suggest that *CDK5* may be a potential regulator in prostate cancer progression through *AR* phosphorylation.

Our study showed that *CDK5* is differentially spliced between prostate cancer and benign tissues at exon 6 (Figure 4B). We calculated isoform expression level of the two isoforms. We found that although *CDK5* was not detected as a DE gene at gene expression level, the relative isoform abundances in prostate cancer and benign tissues are very different. Isoform1 makes up more than 90% of the total expression of *CDK5* in prostate cancer but less than 40% in benign tissues (Figure 4C, Additional file 9). It has been reported that protein *CDK5* encoded by Isoform1 can activate and stabilize *AR* in prostate cancer cell line [44]. *AR* also shows higher expression in the prostate cancer group, with log2 fold change of ~1.43 comparing to the benign group (Additional file 2). We infer that the higher relative expression of Isoform1 in cancer cells plays a role in *AR*-involved prostate cancer progression.

**Figure 4.**
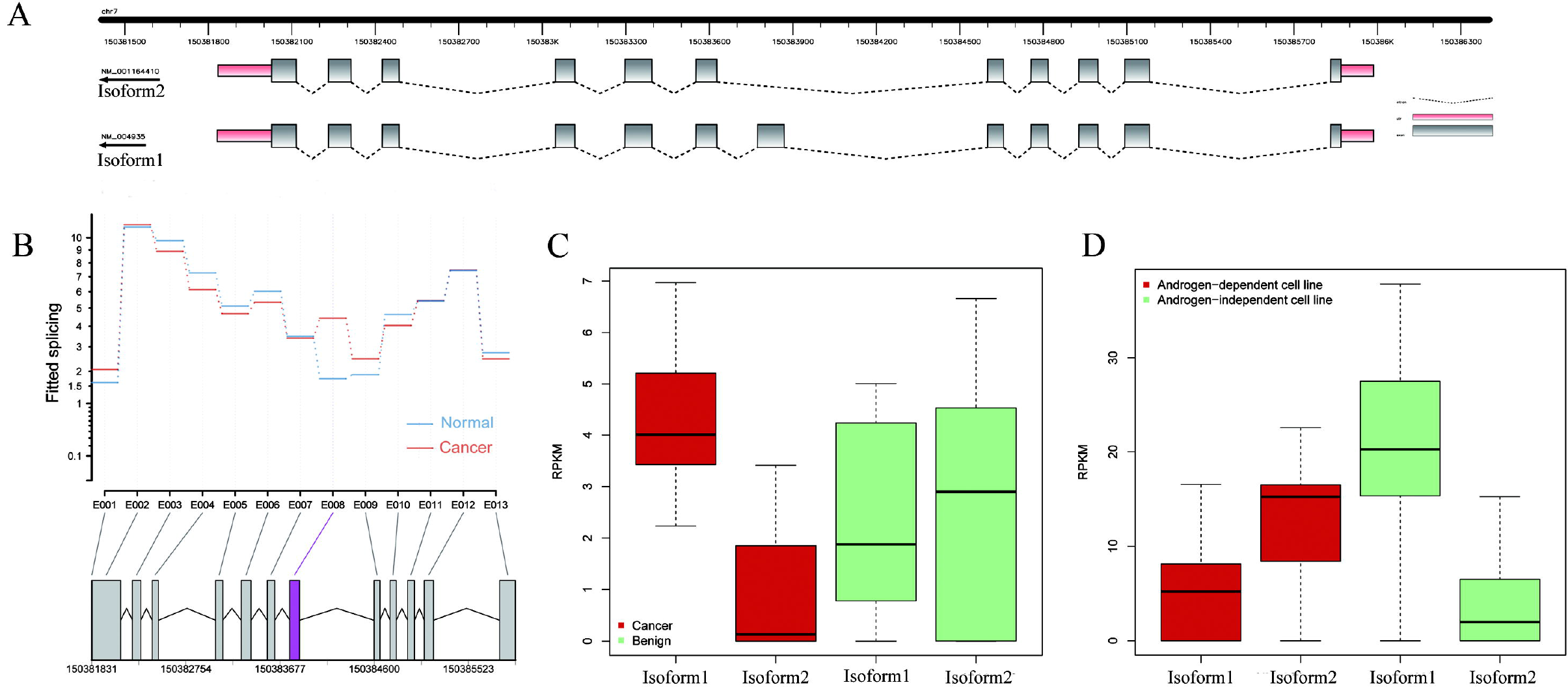
Differential splicing and isoform switching of kinase gene *CDK5* in prostate cancer. (A) The transcript structure of *CDK5* plotted by FancyGene [45]. *CDK5* is located on Chromosome 7 and has two annotated isoforms named NM_004935 (Isoform1) and NM_001164410 (Isoform2). Isoform1 has 12 exons, and its 6th exon is skipped in Isoform2. (B) Differential splicing of *CDK5* illustrated by DEXSeq. The relative exon usage, as measured by “fitted splicing”, was plotted for each exon in two groups. Blue lines are for prostate benign samples and red lines are for prostate cancer samples. The panel at the bottom shows the location of the exons, with the alternative exon highlighted in red. (C) Expression boxplots of the two isoforms in prostate cancer and benign tissue samples. (D) Expression boxplots of the two isoforms in the androgen-dependent and androgen-independent prostate cancer cell lines.

We validated the *CDK5* isoform expression pattern in another RNA-Seq dataset of 21 prostate cancer and benign cell lines published by John Prensner et al. [15].
Significant isoform-switching has not been seen between prostate cancer and benign cell lines. But the benign cell lines still have a lower Isoform1 expression. We found that isoform1 tends to be not expressed or lowly expressed in some prostate cancer cell lines like VCaP and LNCaP, which are androgen-dependent prostate cancer cell lines. We applied clustering analysis on the cell lines by their gene expression profiles, and clustered them into two groups:androgen-dependent cell lines and androgen-independent cell lines. We observed that *CDK5* has different isoform usage between androgen-dependent and androgen-independent prostate cancer cell lines. Isoform1 takes ~28% of total expression in the androgen-dependent group and ~81% in the androgen-independent group, and the proportions of Isoform2 are ~72% and 19%, respectively (Figure 4D, Additional file 9). These results showed that *CDK5* has different isoform preferences in prostate benign, androgen-dependent, and androgen-independent prostate cancer cell lines. Considering that *CDK5* Isoform1 can phosphorylate *AR* at its Ser-81 site and thus stabilize *AR* proteins and *AR* activation to promote prostate cancer cell growth, different usage of *CDK5* isoforms in different prostate cancer development stage can be a modulator of *AR* transcriptional activity.

## Discussion

In this study, we characterized the gene expression profile of prostate cancer by a systematic comparison of RNA-Seq data of prostate cancer and benign tissues. Quantitative analysis was applied on both the gene expression levels and their alternative isoforms, with a special emphasis on kinase genes. The results showed that alternative splicing adds another layer of regulation to gene expression by the different usage of splicing isoforms or different relative expression of alternative isoforms in cancer. Functional enrichment analysis also showed that the differentially expressed and spliced genes are enriched in GO terms and KEGG pathways that are closely related to prostate cancer. Some kinase genes that can phosphorylate *AR* are differentially spliced but none of them are differentially expressed. Differentially spliced exons of detected kinase genes are highly enriched in the protein kinase domain. This indicates that the differentially spliced kinases have the potential to alter protein phosphorylation activity by changing abundance of isoforms with kinase domain.

The study showed that the *AR* phosphorylation kinase gene *CDK5* undergoes isoform switching between prostate cancer and benign tissues. This result was also validated by another RNA-Seq dataset of prostate cancer cell lines. Isoform ratio differences were also observed in androgen-dependent and androgen-independent prostate cancer cell lines. The isoform usage preference of *CDK5* in different prostate cancer development stage may play an important role in prostate cancer cell growth by modulating *AR* transcriptional activity though *AR* phosphorylation and *AR* protein stabilization (Figure 5). These results demonstrated that the isoform ratio change of *CDK5* with different prostate cancer developmental stage may fine tune *AR* activity and contribute to prostate cancer progression. Further research can be carried out to study the influence of isoform ratio change of gene *CDK5* on prostate cancer progression through considering *AR* protein expression and phosphorylation level.

**Figure 5.**
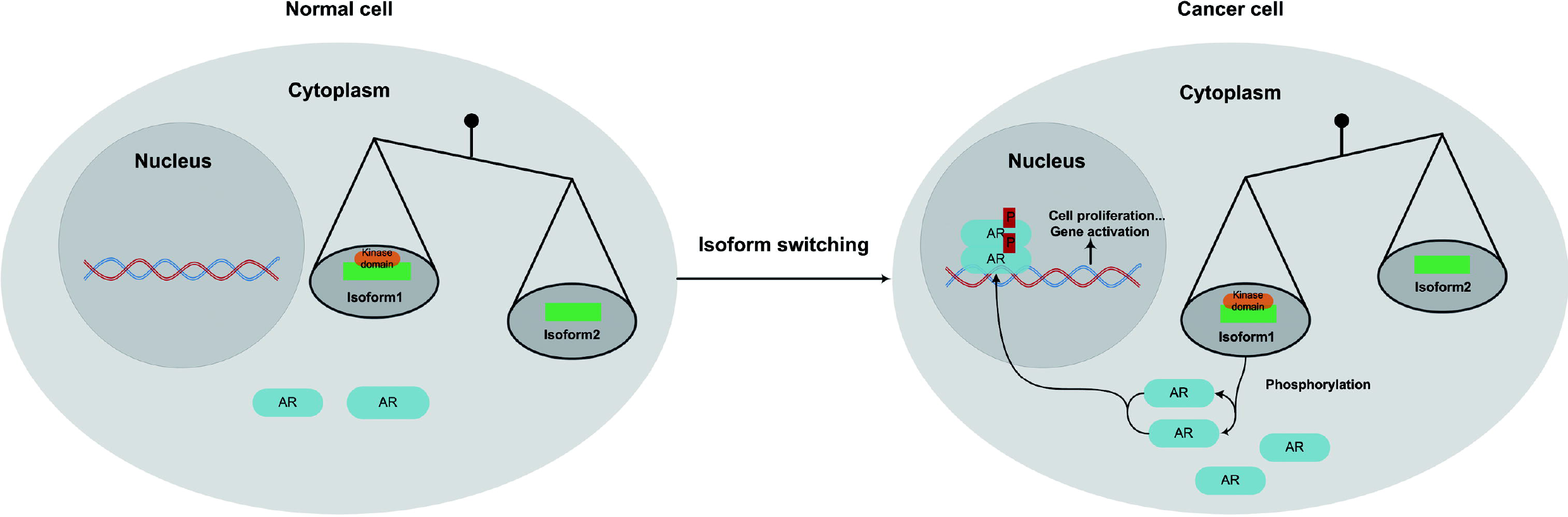
A hypothetic model of the function of *CDK5* isoform switching in prostate cancer. The dominance of Isoform1 over Isoform2 in cancer increases the phosphorylation of *AR*, which increases its protein stabilization and activity in nuclear, and thus promote cancer-related cellular process like cell proliferation.

## Conclusions

In summary, our study provided a transcriptome profile of prostate cancer using RNA-Seq data with an emphasis on differentially spliced kinase genes. It indicated alternative splicing has critical impact on kinase activity in cancers. Especially, isoform switching of kinase gene *CDK5* was found in prostate cancer and benign tissues, which suggests its regulatory role in androgen receptor (*AR*) phosphorylation via alternative splicing. The work brings new understanding to the role of alternatively spliced kinases in prostate cancer and provides an example on the systematic analysis of RNA-Seq data in cancer.

## Authors’ contributions

XZ and TL initiated this work. HF and TL designed the study. HF performed the data analysis under the supervision of XZ and TL. HF and XZ wrote the manuscript. All authors have reviewed and approved the manuscript.

## Acknowledgments

We thank Drs. Likun Wang and Zhixing Feng for insightful discussions. This work is partially supported by the National Basic Research Program of China (2012CB316504), the Hi-tech Research and Development Program of China (2012AA020401) and NSFC Grants 91010016 and 61021063.

## Additional files

### Additional file 1 – Figure S1-S5

Figure S1. DE gene counts in prostate cancer related GO terms. Figure S2. Enriched KEGG pathways by DE genes. Figure S3. REVIGO treemap for enriched GO terms by DS genes. Figure S4. REVIGO treemap for enriched GO terms by DE and DS kinase genes. Figure S5. Kinase domain and phosphorylation site of CDK5 two protein isoforms.

### Additional file 2 – Table S1

DE genes with p-value < 0.05 and FDR < 0.01

### Additional file 3 – Table S2

DE kinase genes with p-value < 0.05 and FDR < 0.01

### Additional file 4 – Table S3

Top 10 up-regulated and down-regulated genes in prostate cancer samples

### Additional file 5 – Table S4

DS genes detected by both DSGseq and DEXseq

### Additional file 6 – Table S5

DS kinase genes detected by both DSGseq and DEXseq

### Additional file 7 – Table S6

GO and KEGG enrichment analysis results for DE and DS genes

### Additional file 8 – Table S7

GO and KEGG enrichment analysis for DE and DS kinase genes

### Additional file 9 – Table S8

CDK5 isoform expression in two datasets

